# “Playing mysterious”: occurrence of Bio-duck calls in Brazil reveals a complex acoustic behaviour for the Antarctic minke whale (*Balaenoptera bonaerensis*)

**DOI:** 10.1101/2021.08.13.456206

**Authors:** Marcos R. Rossi-Santos, Diego Filun, William Soares-Filho, Alexandre D. Paro, Leonardo L. Wedekin

## Abstract

Acoustic methods can provide important data on the occurrence and distribution of migratory species. Information about Antarctic Minke whale (*Balaenoptera bonaerensis*) occurrence in the winter breeding grounds is scarce, mostly limited to old records from whaling stations before 1960’s international moratory, such as Costinha Station in Northeastern Brazil (6° S / 34° W). This work describes the occurrence of the Antarctic minke whale (AMW) through Bio-duck acoustic detections in the Santos Basin, South-Southeastern Brazil (22° and 28° S / 42° and 48° W), registered between November 12 and December 19, 2015. AMW calls were detected for 12 days. We detected and classified 9 different Bio-duck calls in Brazilian coast waters, evidencing a high diverse acoustic behaviour for the minke whale breeding ground. This is the first study to describe the acoustic diversity of AMW vocalizations in lower latitudes, constituting important information to the conservation and management of cetaceans and their habitat. Therefore, our study presents the foremost acoustic evidence of the Antarctic minke whale in Brazil, utilizing high technological passive acoustic methods, such as autonomous underwater vehicle (SeaGlider) sampling.

## Introduction

Marine mammals are key species in the marine ecosystem, generally belonging to high levels in the food chain, controlling natural populations, cycling nutrients, and providing food when decomposing at the oceanic bottom [1]. For some cetaceans, such as the baleen whales, acoustic calls play an important role during foraging and breeding behaviours and have been extensively studied for some species such as the humpback whale (*Megaptera novaeangliae*) [2, 3], the fin whale (*Balaenoptera physalus*) [4, 5], and the blue whale (*Balaenoptera musculus*) [6, 7], although for some whale species the acoustic ecology remains scarcely understood. Furthermore, whale vocalisations can show patterns of occurrence, breeding behaviour, movement, and seasonality of a species within a certain area [8–11] and even geographic differences in acoustic repertoire between different areas [12–15].

The Antarctic minke whale (*Balaenoptera bonaerensis*) (AMW) occurs in the southern hemisphere from Antarctica to near the equator (10° S), performing seasonal offshore migration from polar to tropical waters, like other whale species [16, 17]. The seasonal distribution and migration patterns of nearly all populations of minke whales are poorly understood [18].

Notwithstanding, information on minke whale distribution in the South Atlantic Ocean are scarce, restricted to the whaling station reports along the 20^th^ century, such as Durban, South Africa (29° 53’ S, 31° 03’ E) from 1968 to 1982 [19] and Costinha, Paraiba state, Northeastern Brazilian coast (6° 53′ S, 34° 52′ W), from 1904 to 1985 [20–23] and more recently from a few visual sighting survey [17, 24–26].

With a great technological advance in the last decades, passive acoustic methods have been considered an efficient non-intrusive method to study and monitor cetacean ecology and occurrence along an ocean basin [18, 27–29]. The AMW Bio-duck sound was first described in 2014 [30], after more than five decades of unknown “mysterious” recordings in the Southern Ocean. Sound patterns vary according to different locations, despite occurring simultaneously in areas such as the eastern Weddell Sea and off Western Australia, during austral winter and spring [28,29].

Whale distinctive sounds have been utilized to describe occurrence and distribution patterns for migratory species, highlighting the annual cycle that result in different acoustic environments [2, 27, 31, 32]. Building a broad understanding of the acoustic ecology from a cetacean community is an important step to contribute with marine conservation policies and, ultimately, benefits the human society.

The goal of this work is to describe the AMW Bio-duck calls in a breeding ground off a tropical area, contributing with information about occurrence of this species in the western South Atlantic Ocean. Therefore, this study presents the first acoustic evidence of the Antarctic minke whale in Brazil, utilizing high technological passive acoustic methods, such as autonomous underwater vehicle (SeaGlider) sampling.

## Methods

### Study area

The Santos Basin is situated off the south and southeast Brazilian coast, between 22° and 28° S and 42° and 48° W, in the western South Atlantic Ocean (Fig 1). This basin presents a wide continental shelf, extending almost 200 km offshore in some locations, crossing a diverse depth gradient, and hosts large oil deposits and fisheries resources [33, 34].

**Fig 1.**
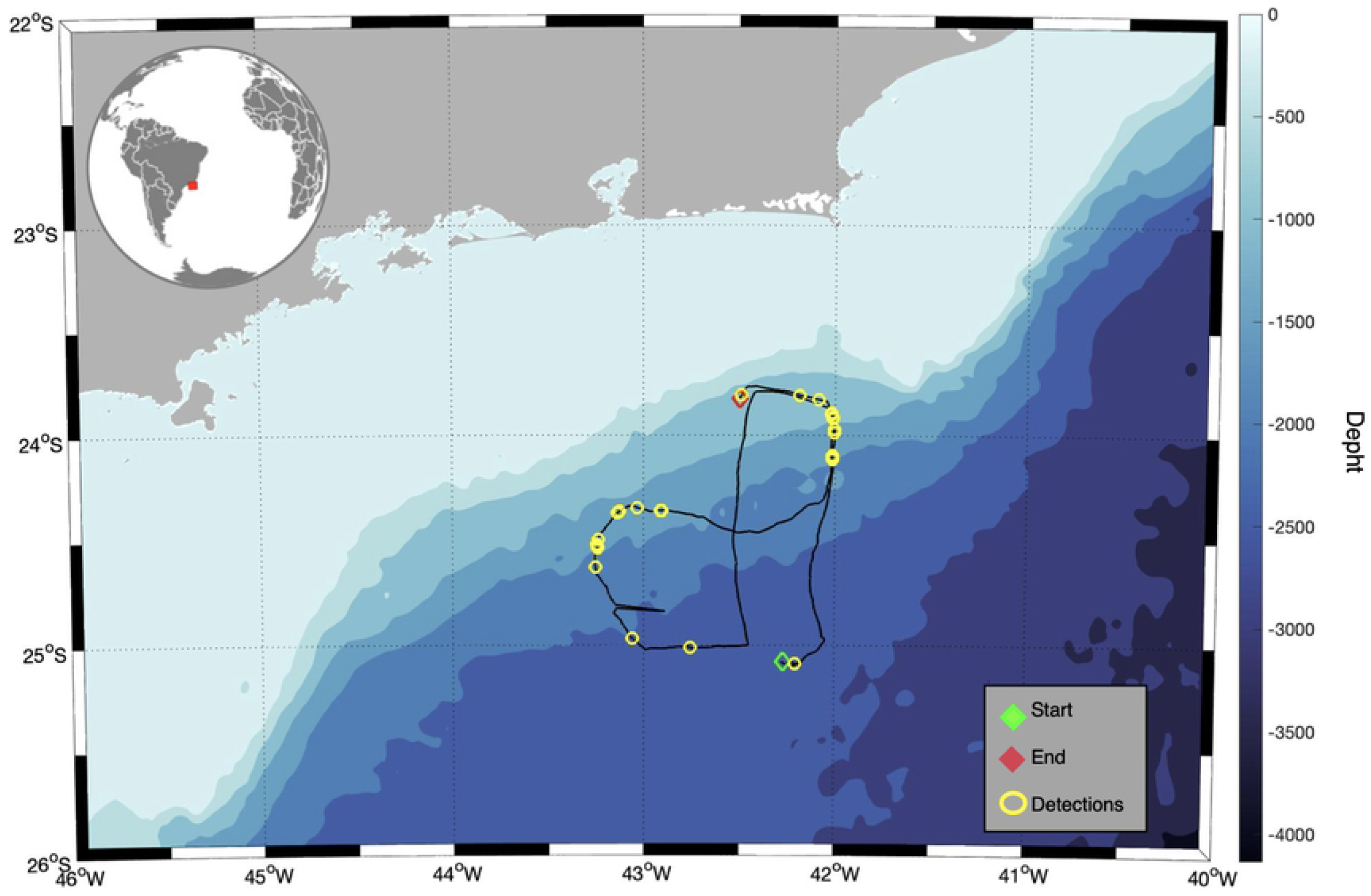
Map of the study area, the Santos Basin, showing the Glider track on Brazilian offshore waters between November 12, 2015 (green marker) until December 19, 2015 (red marker). Yellow dots show the positions where Antarctic Minke Whales Bio-duck calls were detected.

### Data collection

Since November 2015, the Underwater Acoustic Soundscape Monitoring Project (PMPAS-BS, in the Portuguese acronym) collects data in the Santos Basin using different types of methodologies, including a glider, equipped with a custom-designed and in-built passive acoustic recording system. The SeaGlider® (Kongsberg Maritime) is designed for continuous, long-term measurements of oceanographic parameters. The SeaGlider used in this study was equipped with a recording system, composed by a hydrophone HT-92-WB, from High Tech Inc., sensitivity: −165 dB re 1V/ μPa, amplified by 25 dB, and recorded at 125 kHz sample rate and 16-bit resolution. Acoustic data is stored in SD cards (up to 500 hours per survey) in the glider and recovered after each survey. The SeaGlider can perform dives until 1 km depth with a speed of 25 cm/s (0.5 knot). For the present study, we utilized data collected between 12 November 2015 and 19 December 2015

### AMW acoustic detection and classification

In our study, a Bio-duck call was a sound constituted by multiple series of downsweep pulses clustered, separated by <1 second [30, 32, 35]. The Bio-duck call never occurred alone and it always occurred in repetitive sequences [36].

To detect the occurrence of AMW Bio-duck calls in Brazilian waters, the PAMGuard whistle and moan detector [37, 38] was applied over the acoustic files. The acoustic data was decimated to 6 kHz and the spectrograms were generated with 1024 FFT points in windows size of 171 milliseconds and an overlap of 95%. To verify presence or absence events of Bio-duck calls in the detector outputs, the data was manually checked with the ‘Seewave’ package [39] of R software [40], generating spectrograms with Fast Fourier Transform (FFT) length of 1125 samples, 95% overlap and Hanning window.

For the classification of the different AMW Bio-duck calls detected in our data set, we extracted different variables with a custom-built routine in R; able to measure i.e., number of pulses (NP), total duration (TD), average pulse duration (AvPD), average inter-pulse interval (AvIPI), duration first pulse (DFP), duration last pulse (DLP) and peak frequency (PF).

The custom-built routine was generated using the R packages ‘Seewave and ‘warbleR’ [41]. The algorithm is an amplitude detector capable to automatically extract and calculate different measurements using the waveform of the signatures of interest. It works detecting and measuring the downsweep signature of the AMW Bio-duck calls between 40-450 Hz. For the analyses in depth, the audio files were decimated to 6000 Hz and a frequency filter between 40-450 Hz with an amplitude filter of 20 dB was applied, making sharpen the Bio-duck pulse spectrum.

The amplitude threshold was calculated according to the signal-to-noise ratio (SNR) of every AMW Bio-duck call detection. The SNR formula used was *SNR* = *20*log10(rms(signal)/rms(noise))*. The noise value was measured during 0.6 seconds in the decimated audio files before the detected call event in the frequency bounds between 40 – 500 Hz and the signal value was measured during 1.5 seconds after the noise value. For acoustic detections with a SNR<5 dB we estimate a threshold=50, with SNR>5 dB and SNR<10 dB we estimate a threshold=30, with SNR>10 dB and SNR<15 dB a threshold=20, with SNR>15 dB and SNR<20 dB a threshold=15 and with an SNR>20 dB a threshold=12 (Fig 2). Because of the differences in quality of call detections (SNR) our algorithm for automatic classification was implemented to use the low frequency downsweep component present in all Bio-duck calls. Every amplitude event detected must be in the time range of 0.055-0.9 seconds, proper range of an AMW Bio-duck pulse (calculated in this study).

**Fig 2.**
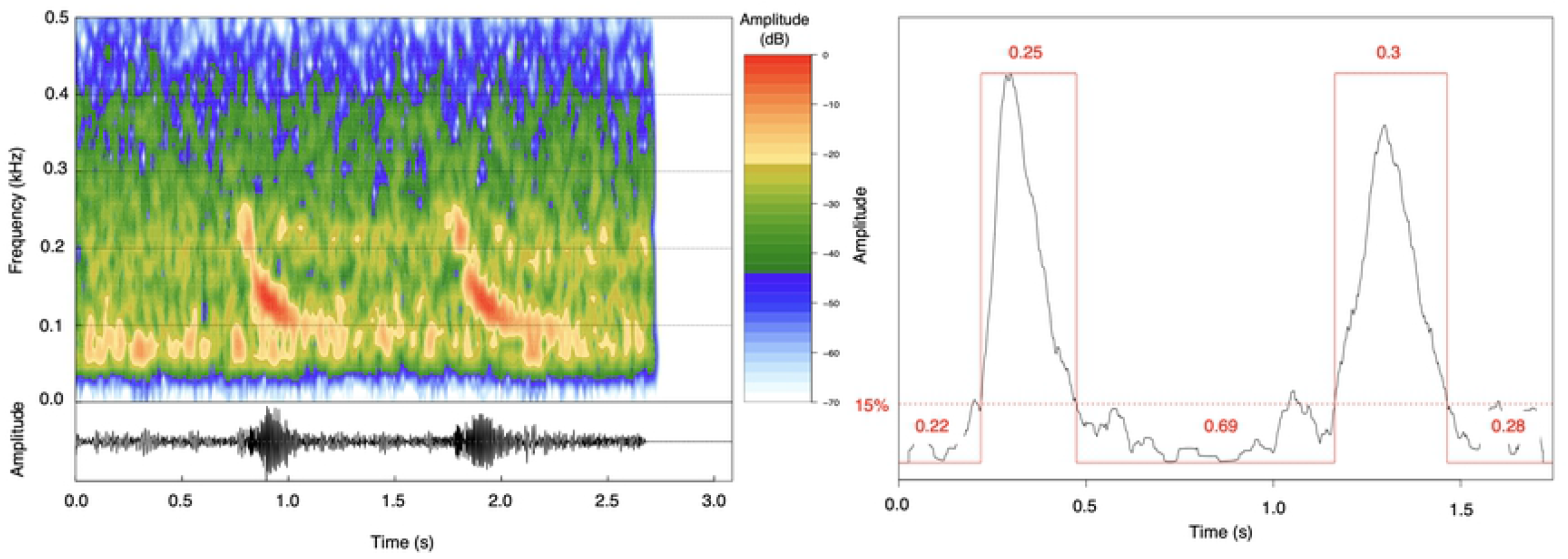
Spectrogram and waveform of the downsweep component of the Antarctic minke whale Bio-duck “B4”. Waveform of the bio-duck “B4” showing the threshold in 12% to identify and measure the different pulses and inter-pulse-intervals (IPIs) in seconds.

We separated the different Bio-duck calls using clustering analysis based on the calculated acoustic parameters. An Euclidean method was used to calculate the distances between the different clusters. To determinate the number of *k* we used the ‘NbClust’ R package [42], which automatically calculates and provides 30 different indices for determining an appropriate number of *k* from the different results obtained by varying all combinations of number of clusters, distance measures and clustering methods.

The different parameters extracted from each Bio-duck call type were used in a hierarchical agglomerative cluster analysis to generate unsupervised call type groups. The information from the matrix of distances calculated a dendrogram to identify the dissimilarities and connections between the different Bio-duck call types detected in Brazilian waters.

## Results

Between November 12 to December 19, 2015, the SeaGlider covered approximately 725 km of distance and collected 262 hours of acoustic data during 187 glider dives. AMW calls were detected along the transect for 12 days Table 1. Manual analyses confirmed the detections were AMW Bio-duck calls, based on the literature description [30, 32, 35, 43].

**Table 1.** Antarctic minke whale Bio-duck call encounters in Brazilian offshore waters.

| Date | Hour start | Hour end | Latitude S° | Longitude W° | Glider depth (m) |
| --- | --- | --- | --- | --- | --- |
| 12.11.2015 | 13:49:00 | 13:54:41 | -23.80815506 | -42.48846817 | 112.98 |
| 14.11.2015 | 08:26:00 | 08:43:40 | -23.80920029 | -42.18441772 | 1.30 |
| 14.11.2015 | 15:47:00 | 16:04:40 | -23.82978058 | -42.08476257 | 279.75 |
| 15.11.2015 | 03:16:00 | 03:49:47 | -23.89512062 | -42.02005386 | 130.18 |
| 15.11.2015 | 03:50:47 | 04:14:46 | -23.89885521 | -42.01930618 | 52.63 |
| 15.11.2015 | 05:46:00 | 06:21:21 | -23.9147644 | -42.01307678 | 345.10 |
| 15.11.2015 | 06:23:21 | 06:57:20 | -23.91983223 | -42.01108932 | 118.74 |
| 15.11.2015 | 11:50:00 | 12:21:07 | -23.97163963 | -42.00485611 | 876.99 |
| 15.11.2015 | 13:31:05 | 13:41:05 | -23.98914909 | -42.0019455 | 108.62 |
| 16.11.2015 | 00:12:00 | 00:43:19 | -24.09622574 | -42.01313782 | 947.00 |
| 16.11.2015 | 00:45:19 | 01:17:18 | -24.10261917 | -42.01353836 | 874.85 |
| 16.11.2015 | 01:19:18 | 01:51:17 | -24.10923958 | -42.01395416 | 645.67 |
| 16.11.2015 | 01:53:17 | 02:17:16 | -24.11581612 | -42.01436615 | 435.60 |
| 22.11.2015 | 07:37:27 | 08:09:26 | -24.35966492 | -42.90554047 | 129.98 |
| 22.11.2015 | 08:11:26 | 08:23:26 | -24.35882759 | -42.90994263 | 177.80 |
| 23.11.2015 | 09:39:00 | 10:08:33 | -24.34605026 | -43.03238297 | 761.20 |
| 24.11.2015 | 13:26:00 | 13:38:07 | -24.36293983 | -43.12480164 | 150.02 |
| 24.11.2015 | 15:16:16 | 15:44:15 | -24.37080193 | -43.13537979 | 121.74 |
| 25.11.2015 | 17:30:52 | 17:52:51 | -24.49822235 | -43.2351265 | 525.01 |
| 25.11.2015 | 21:12:00 | 21:43:14 | -24.52141762 | -43.24263382 | 1.28 |
| 25.11.2015 | 23:13:16 | 23:43:15 | -24.53253174 | -43.2402153 | 589.22 |
| 25.11.2015 | 23:45:15 | 23:50:15 | -24.5354538 | -43.23957825 | 198.02 |
| 26.11.2015 | 00:01:15 | 00:11:14 | -24.53692055 | -43.23926163 | 6.73 |
| 26.11.2015 | 15:11:26 | 02:39:00 | -24.63012505 | -43.2525177 | 133.27 |
| 29.11.2015 | 22:37:30 | 23:44:49 | -24.96931648 | -43.05936813 | 409.64 |
| 01.12.2015 | 19:34:00 | 19:41:03 | -25.01185799 | -42.75591278 | 546.45 |
| 17.12.2015 | 14:27:00 | 14:44:00 | -25.08859062 | -42.20511627 | 4.25 |

### Description of vocalizations

The vocal classes or groups were named according to acoustic properties extracted automatically of Bio-duck calls detected in Brazilian waters based on automatic classification Table 2 and indicated in capital letters followed by a number. This number value corresponds to the number of down swept pulses present in a Bio-duck call classified (Fig 2). Call types were qualitatively named, to maintain consistency between naming schemes and sound structure. For the automatic measurements we did not consider the harmonics because they varied in relation of the proximity of the animals to the SeaGlider. Even though, we show spectrograms with the full frequency range to distinguish the harmonics in the Bio-duck calls detected up to 2000 Hz.

**Table 2.** Antarctic Minke Whale bio-duck call extracted measurements with the amplitude detector.

| Cluster | N° Pulses | Total Duration (s) | Duration Av pulse (s) | Duration Av IPI (s) | Duration 1st Pulse (s) | Duration Last Pulse (s) | Peak Freq (Hz) | (n) |
| --- | --- | --- | --- | --- | --- | --- | --- | --- |
| A | 2 | 1.26 | 0.28 | 0.7 | 0.28 | 0.29 | 131.75 | 30 |
| A | 3 | 1.3 | 0.31 | 0.19 | 0.32 | 0.3 | 182.97 | 30 |
| A | 4 | 1.61 | 0.28 | 0.17 | 0.33 | 0.1 | 177.36 | 30 |
| B | 3 | 1.04 | 0.09 | 0.38 | 0.08 | 0.09 | 162.91 | 14 |
| B | 4 | 1.45 | 0.19 | 0.23 | 0.18 | 0.09 | 181.64 | 30 |
| B | 5 | 1.7 | 0.23 | 0.14 | 0.27 | 0.1 | 159.84 | 30 |
| C | 10 | 1.45 | 0.08 | 0.05 | 0.07 | 0.08 | 245.58 | 30 |
| D | 6 | 1.39 | 0.14 | 0.11 | 0.11 | 0.11 | 116.25 | 30 |
| D | 7 | 1.62 | 0.16 | 0.08 | 0.18 | 0.06 | 86.31 | 30 |

### Bio-duck calls classification

Nine different Bio-duck calls (Fig 3) were detected and classified with a hierarchical agglomerative cluster analysis (Fig 4). Every call analyzed contained between 2 and 11 pulses. The duration of a Bio-duck sound was between 1.26 – 1.70 seconds and the average pulse duration was between 0.05 – 0.38 seconds. The AvIPI was 0.05-0.38 seconds. For all categories, whenever a Bio-duck call incremented the number of pulses, the AvIPI decreased while the total duration increased. The duration of the AvFP varied from 0.07 – 0.33 seconds and the last pulse was between 0.06 – 0.3 seconds. The peak frequency of the down swept component was 86.31-245.48 Hz Table 2. The analysis identified 4 different clusters (73%), grouping the “A2”, “A3” and “A4” calls in the cluster A, Bio-ducks “B3”, “B4” and “B5” in the cluster B, and only one call, “C10”, in the cluster C. Finally, the cluster D was composed by two calls: “D6” and “D7” (Fig 5).

**Fig 3.**
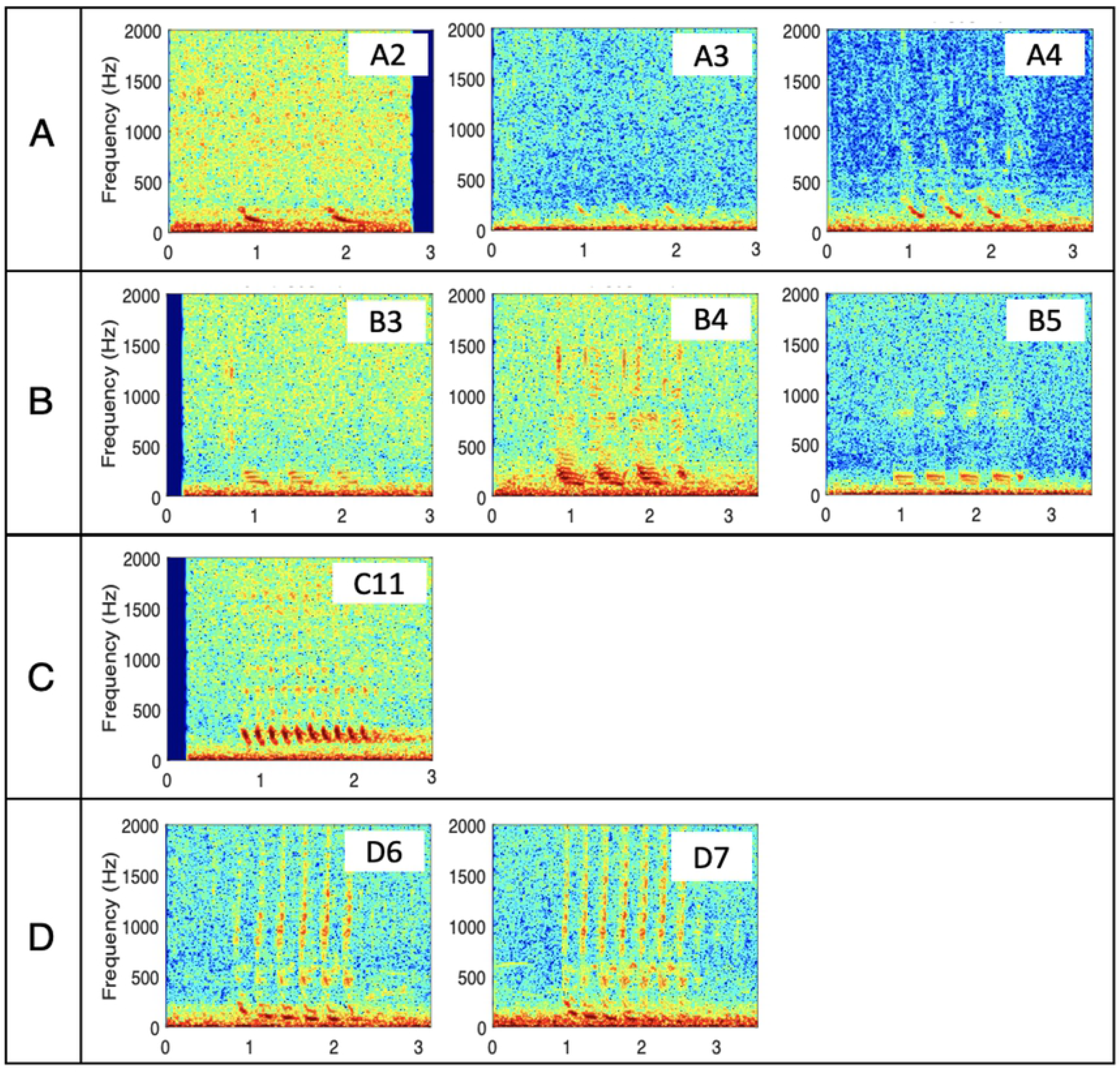
Spectrograms of the different Bio-duck calls of the Antarctic Minke Whale detected in the Santos Basin, Brazilian offshore waters. FFT=512, overlap= 95%, Time expressed in seconds and frequency scale in Hz.

**Fig 4.**
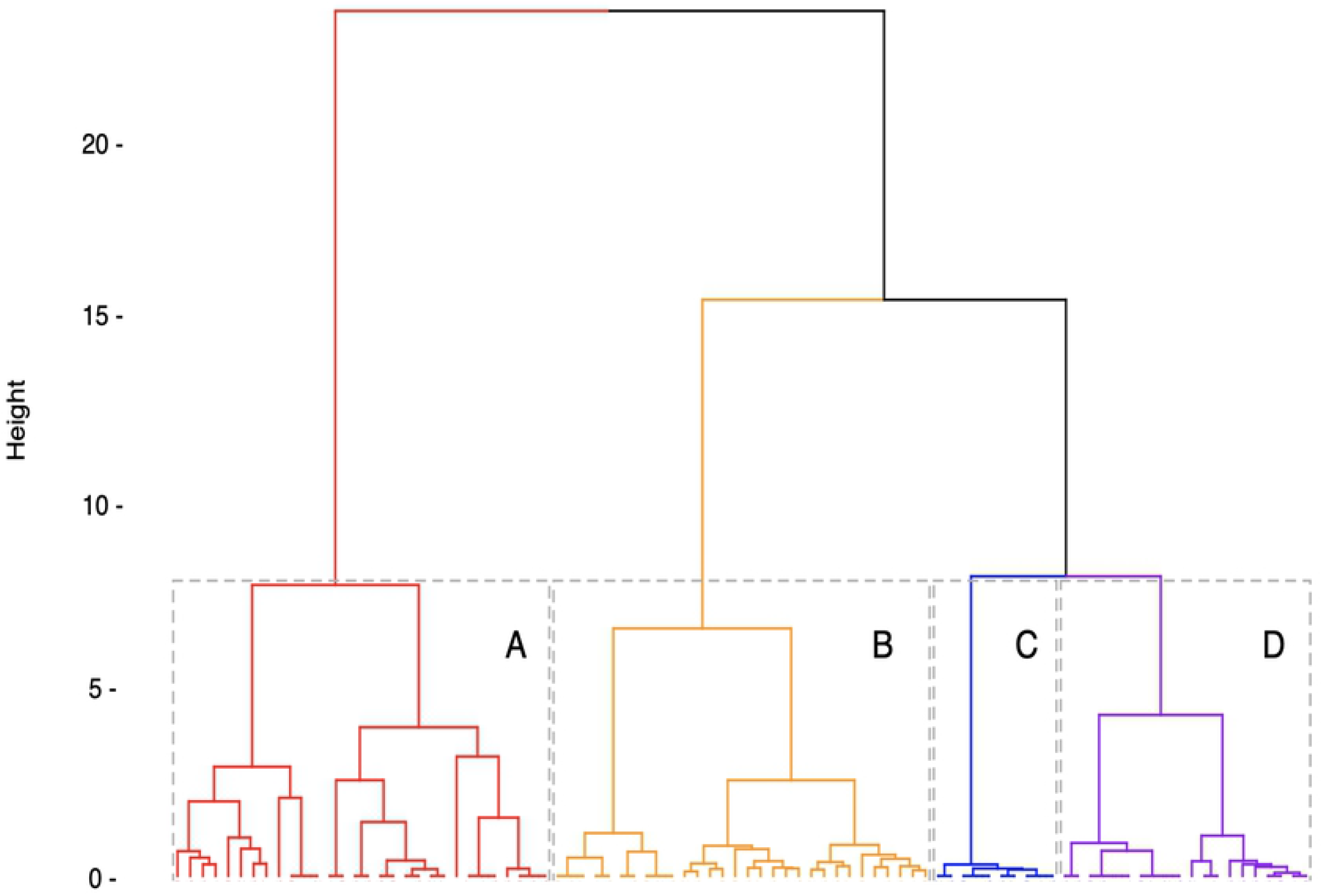
Dendrogram showing results of a hierarchical cluster analysis. Clusters representing the four classes of Antarctic minke whale Bio-duck calls are identified numerically and with brackets (color online). Cluster A corresponds A2, A3 and A4; Cluster B with the B3, B4 and B5 sounds; Cluster C with the C11; Cluster D with D6 and D7 calls.

### Bio-duck call occurrence

The Bio-duck call named “D6” was the most common call detected in the data, being present during 7 different days. The second most common call detected was the Bio-duck “A4” presented in 4 days. The Bio-duck “B4” was detected in 3 different days. The others Bio-duck calls were detected only one day along the glider deployment period analyzed. During the 15/11/2015 three different Bio-duck calls were detected. It was the day with most acoustic diversity of Bio-duck call occurrence, from 3:16 am until 1:41 pm. During that day, the Bio-duck calls “D6”, “A4” and “C11” were detected. The second day with most acoustic variability was the 25/11/2015 with the presence of two different call types (“D6” and “A2”). During the rest of the days with acoustic presence, the type of call detected was repeated throughout the day Table 1.

## Discussion

This work evidenced the presence of 9 different Bio-duck calls registered for the Brazilian coast, a low latitude presumed breeding ground for migratory baleen whales in the western South Atlantic Ocean. This expresses the highest diversity of acoustic behaviour reported for the species so far. The diversity of calls found suggests that the AMW may produce organized and repetitive sounds, which suits in the description of songs for other whale species, such as the humpback whale [2], southern right whales [44] and north Atlantic right whales [45]. In addition to the variability of sounds produced by AMWs, their vocalizations are produced in repetitive sequences just like those of, for example, Antarctic blue whales [46] or fin whales [47–49].

### Bio-duck calls comparison

Regarding acoustic structure, the Bio-duck has been considered a very conspicuous signature call [18, 30, 36, 43]. The most obvious characteristic is that energy is in a frequency band from 50 to 500 Hz [30, 32, 35, 43], although more intense signals harmonics have been observed up to 1 kHz.

The first study attributing the Bio-duck sounds to AMWs described the calls in series between 5-12 pulses, produced in regular sequences with an inter-sequence interval of 3.1 s. The data were collected from a tagged AMW in Wilhelmina Bay, in the Antarctic Peninsula, and they described 3 different Bio-ducks composed between 3-7 pulses and 3 types of downsweep low-frequency [30]. Another study in the Antarctic Peninsula [32], using year round acoustic data from a mooring position, described 4 distinct Bio-duck call variants, with one variant having two sub-types. The calls were described with sequences between 4-13 pulses. In addition, this study also indicates the presence of a call type of downsweep low-frequency. Recently, [35] described the occurrence of AMW calls in South African waters and Maud Rise, Antarctica. The vocalisations detected in the study presented harmonics up to 2000 Hz and were classified in 3 categories with one category presenting 2 sub-types. The Bio-ducks presented sequences between 4-10 pulses. In Perth Canyon, western Australia, [43] used a time ratio method for detecting the bio-duck signal, distinguishing two different types of call. One with a low repetition rate (T=1.6 seconds) and one with approximately doubled period (T=3.1 seconds).

The calls described in our study match some of the Bio-ducks previously described in Perth Canyon [43], Western Antarctic Peninsula [30, 32], South Africa and Maud Rise, Antarctica [35]. Similarly, 7 of 9 calls detected in Brazil, classified in different categories, presented harmonics up to 2000 Hz. All the calls described for South Africa were present in Brazil. The only difference between the two localities is that the C-type vocalisation in South Africa was composed of 4 pulses, while in Brazil it was recorded with 6 and 7 pulses. This information evidences the complex structure of the Bio-duck calls, which can be found in general categories, but also presenting subtypes of sounds, according to the different number of pulses. With the information that AMW Bio-duck calls can be classified into categories, studying the subtypes of sounds could help to better understand the acoustic behaviour of this species and even know if through acoustics it is possible to identify populations.

### Acoustic seasonality

Comparing the present data with other regions, the acoustic seasonality seems to match. In Brazil (24 and 25° S), we recorded Bio-duck calls during November and December, while collected between October and December. Likewise, Bio-duck calls were recorded in the Perth Canyon (32° S) [43]. In Namibia (20° S), a double peak was recorded between June-August and November-December [50]. In South African waters (34°S) a peak of acoustic activity was described between September-October [35] and in the Juan Fernandez Archipelago, off Chile, (33°S) a year-round acoustic presence was recorded with a peak between May-August [51].

When comparing low and high latitude areas, there appear to be similarities in the acoustic seasonality of AMWs. In the Southern Ocean, specifically in the western Antarctic Peninsula, periods of acoustic activity have been described for AMWs between the months of May-November with peaks during July-October [32]. A similar situation occurs in the Weddell Sea, where the acoustic presence has been described between May-December, with a peak during June-November [28, 32, 36]. The similarity in the seasonality of the acoustic behaviour of AMWs between low and high latitudes indicates that not all the individuals migrate to the breeding ground. Part of the population possibly stays in Southern Ocean waters for a longer time than in high latitudes. The fact that AMWs exhibit simultaneous acoustic behaviors in different regions also shows that there are probably distinct populations. Moreover, the fact that the acoustic behaviour occurs simultaneously in high and low latitudes infers that the sound emission is related to animal behaviour rather than to environmental conditions.

### Antarctic minke whales in Brazil

Few is known about the distribution and ecological relations of baleen whales from the genus *Balaenoptera* in their breeding grounds (IWC, 2021). For the Antarctic feeding ground, it is known that despite an evident habitat segregation between minkes and humpbacks they can overlap their distribution close to the edge-ice [52, 53]. Regarding the AMW distribution, our recordings in Santos Basin were made between 24 and 25° S, while the Perth Canyon lies at 32° S. This geographic area could represent a different temporal niche choice, comparing to other frequent whale species such as the humpback whale. Similarly, the presence of AMWs, through the bio-duck calls, indicate that this species is occurring on the Brazilian continental slope and oceanic waters, as suggested by other recent observations [17, 24–26], possibly in a search for an emptier acoustic habitat, once humpback songs predominate in the soundscape of the continental shelf during breeding seasons.

We registered the presence of AMW in November and December in a low latitude area, such as the Brazilian coast. The species also occurs in their feeding areas in high latitudes of Antarctica in this period [36]. This information raises consideration that is possible that part of the AMW does no migrate, staying in the low latitude areas, where they could find enough resources to live year-round, as occur with other species/populations, such as the humpback whale from the Arabian sea population [54, 55].

## Conclusion

In summary, our results show that Antarctic minke whales perform a diverse repertoire of Bio-duck calls in oceanic waters off Brazil. The distinctiveness of these type of calls in Santos Basin reinforce the occurrence of these species in the region, that is further south than the main areas of whaling, where their presence is well documented through historical accounts. Further studies using passive acoustics to detect Bio-duck calls may raise important information about this species seasonality, migratory timing and connections with other feeding and breeding regions of the southern hemisphere

Once the Bio-duck calls were registered in deeper waters, it seems that minke whale distribution does not completely overlap with the distribution of humpback whales in the western South Atlantic breeding ground. This pattern could be related to a certain spatial acoustic niche, as the humpback whale sounds predominate in the soundscape during their breeding season [56].

This study also highlights the importance of further implementation of passive acoustic monitoring in Brazil, nowadays limited to a few initiatives as requirements for oil and gas seismic and exploitation activities. We suggest the use of passive acoustic monitoring should be a mandatory routine to study cetacean occurrence, behaviour, and potential noise impacts on them, in many other marine exploitation activities, such as port constructions and cargo ship traffic, considered to impact Brazilian coast without any previous assessment [57]. Further investigation with a broader acoustic dataset can elucidate important questions about the minke whale annual cycle, such as the migration timing, if they are breeding, foraging or doing both vital activities while in their tropical breeding grounds.

## Acknowledgments

We thank Petrobras for kindly authorizing the data use from Projeto de Monitoramento da Paisagem Acústica na Bacia de Santos (PMPAS-BS). This Project is part of a monitoring program required by Brazil’s federal environmental agency, IBAMA, for the environmental licensing process of the oil production and transport by Petrobras at the Santos Basin pre-salt province. We thank Denise Risch and Ilse Van Opzeeland for putting the conceptual authors in initial contact, resulting in this manuscript.

## Author Contributions

MRRS and DF have equal contributions as leading authors of this study. Conceptualization: MRRS DF. Data curation: WSF ADP MRRS LLW. Investigation: MRRS DF WSF ADP LLW. Methodology: WSF MRRS DF. Formal analysis: WSF ADP MRRS DF. Data Validation: WSF ADP MRRS DF. Contributed analysis tools/software: WSF DF. Project Management: LLW. Writing – original draft: MRRS DF. Writing – review & editing: MRRS DF WSF LLW.

